# A conditional generative model to disentangle morphological variation from batch effects in model organism imaging studies

**DOI:** 10.1101/2025.06.04.657966

**Authors:** Ricardo M. Valdarrago, Hongru Hu, Ruoxin Li, José Miguel Uribe-Salazar, Megan Y. Dennis, Gerald Quon

## Abstract

Identifying organism-level phenotypic variation that arises from genetic variation is a longstanding problem in genetics. Model organisms such as zebrafish are frequently used for genotype to phenotype studies as they can be genetically manipulated, grown and phenotyped for morphological changes in a high throughput manner through automated systems and imaging. However, individuals from model organisms are typically grown in groups: clutches for zebrafish and frogs, and litter for rodents. These clutches act as strong confounders during image analysis, as individuals from different genotypes but grown in the same clutch tend to look more similar to each other than to individuals with the same genotype in other clutches. Existing approaches such as conditional image classification models and domain adaptation approaches perform poorly on addressing these technical batch effects. Here, we propose a conditional latent diffusion model (cLDM) that disentangles these technical batch effects from morphological features by explicitly conditioning on batch-specific variables during generation, enabling targeted separation of technical artifacts from biologically relevant data. This approach enables accurate classification of genotypes using morphological images of individual zebrafish from distinct mutant classes. Furthermore, this approach allows us to efficiently correct for batch effect and demonstrates the versatility of cLDM in tackling domain-specific problems. This work highlights the potential of cLDM to overcome batch effects and extract meaningful features.

## INTRODUCTION

Model organisms such as the mouse and zebrafish are instrumental in studying gene function due to their genetic similarity to humans, ease of genetic manipulation and in the case of zebrafish, their transparent embryos facilitate imaging^1^. In recent years, detecting phenotypes in larval zebrafish has involved imaging-based analysis of morphological features^2^, including the development of high-throughput phenotyping protocols to identify gene knockouts with similar phenotypes^3^. In a typical case-control genetic experiment in zebrafish for example, fertilized embryos from the same clutch (group of embryos from the same parents, generated at the same time) are divided into three groups: an injected group that receives CRISPR/Cas9 or morpholino targeting of a specific gene, and two control groups: an uninjected control group, and a mock-injected control group that is injected with a scrambled gRNA. Both the injected and control embryos are grown on the same 96-well plate to minimize batch (clutch) effects, then imaged and genotyped for genotype to phenotype analyses. During analysis, the injected for one genotype can be compared to the controls from the same clutch to identify significant changes in morphology, compared to controls^4^

However, because each injected is compared to a control from the same clutch and plate to minimize batch effects, and given that each plate can only support up to 96 individual embryos, typical studies must distribute genotypes (and their respective injected individuals) across multiple plates. Due to batch effects across plates and clutches that cause zebrafish from the same clutch and plate to possibly be more morphologically similar than to those from other clutches, it is challenging to compare two different mutant genotypes to determine to what extent their associated morphological changes are consistent.

Removing systematic differences between sets of (genotype) labeled images is commonly formulated as a domain adaptation (DA) problem, in which clutches or batches are treated as domains^5^, and the goal is to ultimately train a classifier using all of the genotypes (labels) present across all of the domains simultaneously. Domain adaptation methods can be divided into different categories based on the assumptions they make about overlap of label sets between different domains. The most common approach is closed set domain adaptation (in which all domains have the same classes/genotypes)^6,7^, the experimental setup described above demands approaches capable of multi-source domain adaptation with partial label overlaps between domains^8,9,10,11,12,13^. Furthermore, in the genotype to phenotype experiments, zebrafish can have multiple types of labels as well (genotypes, age, clutch) which makes it difficult using existing approaches to handle different types of labels simultaneously during training.

Here we propose a conditional latent diffusion model (cLDM) to address this domain shift problem. In the reverse process of this cLDM, we condition generation of the image on the multiple label sets (genotype, batch, age) and train across all genotypes, batches and ages simultaneously in order to disentangle the effects of the three label sets. Our cLDM simultaneously uses the clutch-specific controls to determine the effect of each mutation, and uses the controls present in every clutch to determine the clutch- and age-specific effects on image (morphology). Applied to imaging data from 3,146 zebrafish larvae representing 17 zebrafish gene mutants as well as controls, our approach allows us to mitigate clutch effect across all zebrafish, and compare gene mutants from different clutches in order to identify groups of mutations which yield similar morphological phenotype, which are not evident from analysis of the original images alone and yield biological insight into how those mutations are functionally related in the zebrafish genome. We also demonstrate our approach performs better at removing clutch effect compared to existing state of the art multi-source domain adaptation methods, suggesting the cLDM is a useful model for multi-source domain adaptation when multiple label sets are present.

## RESULTS

### cLDM framework and analysis overview

The cLDM model has two components: a Variational Autoencoder (VAE) to project images to a low-dimensional embedding that is more efficient and less noisy compared to the original images, and a Diffusion Model (DM) for learning how the label sets drive variation in VAE image embedding space (**Figure 1a**). The DM is composed of two parts: a forward and reverse process. The forward process adds small increments of Gaussian noise over many steps to transform the distribution of images in the VAE embedding space into an approximately isotropic multivariate Gaussian distribution (**Figure 1b**). Input image embeddings are then passed through a reverse conditional denoising process that learns to iteratively denoise a sample drawn from an isotropic Gaussian distribution, to yield samples that approximate the input training data distribution. Our reverse denoising process is conditioned on the label sets of the image (clutch, age and genotype of the zebrafish), so that during training across many clutches, ages and genotypes, the model learns to disentangle the effects of the different label sets.

**Figure 1:**
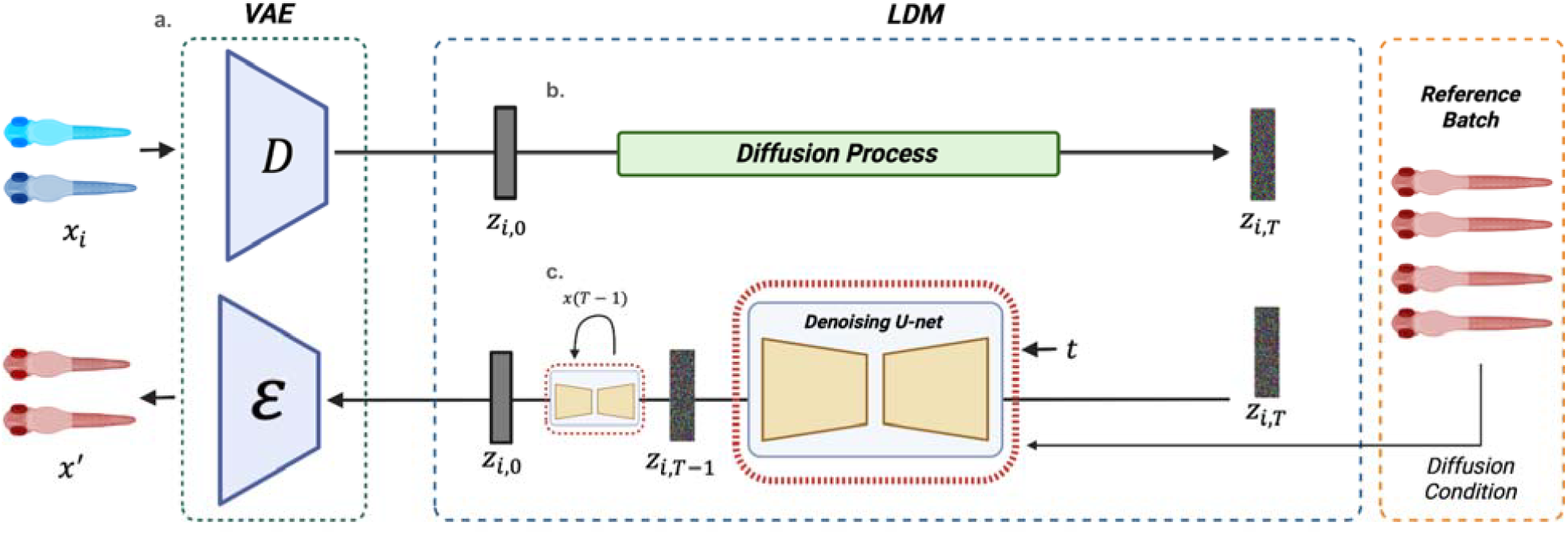
Conditional latent diffusion model (cLDM) overview. **(a)** A variational autoencoder (VAE) is used to project imaging data to and from an embedding space in which the latent diffusion model operates. **(b)** Forward diffusion process iteratively adds noise for *T* time steps to the latent representation of *z* producing a fully noised *z*_*T*_. **(c)** *z*_*T*_ is passed through the denoising reverse process that iteratively denoises the embedding until reaching *z*. During this process an additional condition term *C* encoding biological and technical variables is applied to guide the generation based on a given condition.

Here we outline the overall strategy for image analysis using cLDM. The goal of our cLDM model is to mitigate the effect of two label sets (clutch, age) in a manner that is independent of the third label set (genotype) so as to prevent label contamination of downstream tasks; identifying effects of genotype on zebrafish morphology occurs as a subsequent image classification task performed on the zebrafish images adjusted for clutch and age effect. The principal reason why we do not perform domain adaptation and classification jointly as is done in some other approaches (CITE) is so we can perform additional, unsupervised learning analysis of the adjusted images to yield biological insight, as demonstrated in our results below.

We first obtained a set of 3,146 zebrafish images from Soto et al (CITE) and standardized them with image preprocessing (**Figure 2a**), and obtained their label sets. Their label sets consist of discrete labels for genotype, clutch and age (see **Methods**). There are 30 genotypes (including three types of control genotypes), two zebrafish age groups (3dpf (days post fertilization) and 5dpf) and 22 clutches. Individuals with each genotype-age combination are profiled in one or more clutches. In order to mitigate the effect of two label sets (clutch, age) independently of the third label set (genotype), for each clutch, we modify the labels of both the mutant genotypes as well as the control genotypes (**Figure 2b**). For the mutant genotypes, we assign them a unique mutant label that is not shared with any other clutch. For the control genotypes, we divide them into two groups: one with the original control genotype label (that will be shared across clutches; the “shared controls”), and the other is given a unique control genotype label that is not shared with any other clutch (“held-out controls”) and is therefore treated equally with the mutant genotypes. By providing these re-labeled images to cLDM to train, we allow cLDM to use the common control genotype labeled images to learn clutch and age effects across batches. As will be discussed next, the relabeled mutants and controls will be used to evaluate performance of both the cLDM and the image classifier.

**Figure 2:**
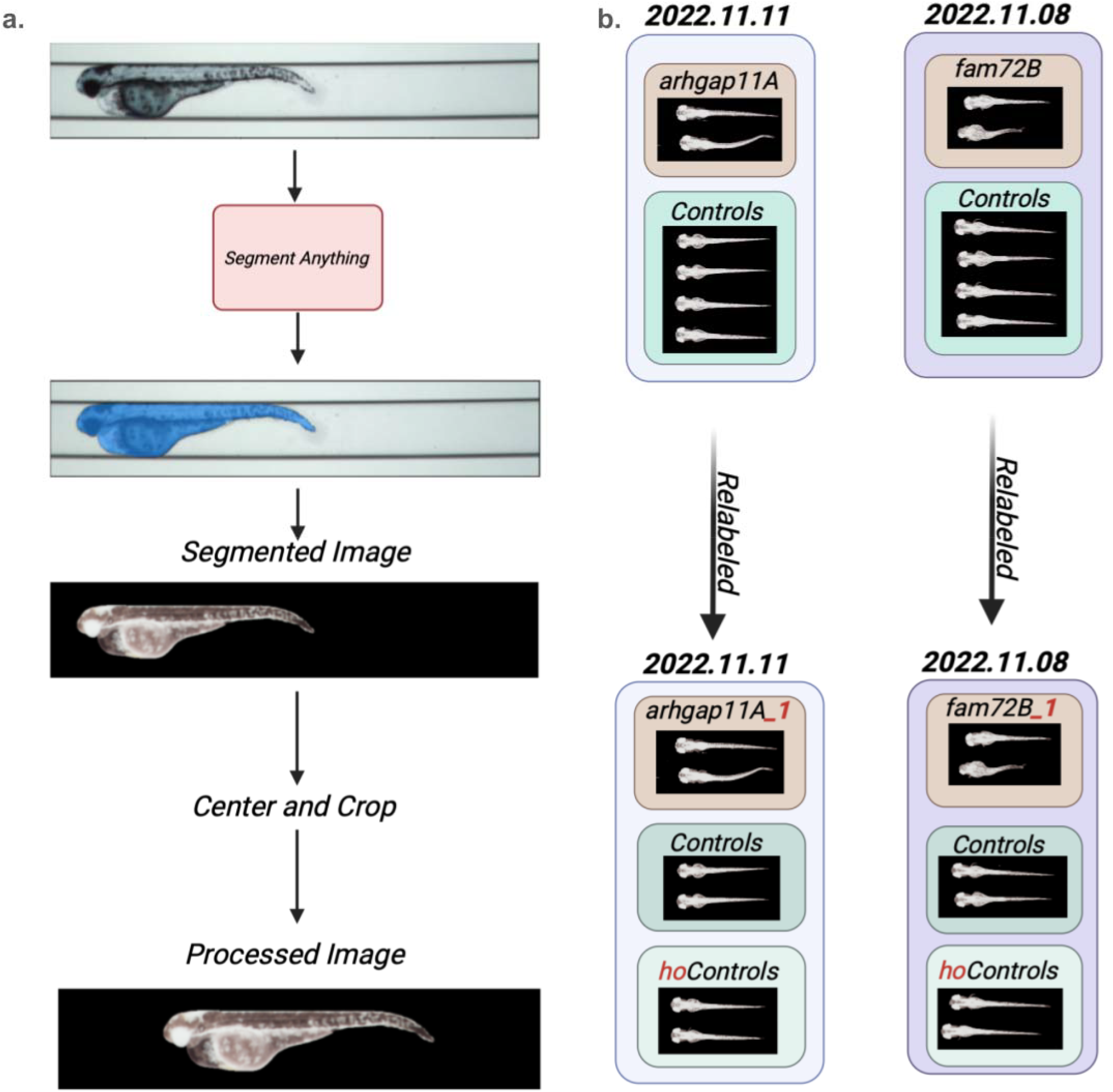
Preprocessing pipeline and dataset relabeling. Our zebrafish preprocessing pipeline starts with using the Segment Anything Model (SAM) to capture segmentation masks and remove the background. We then subsequently perform a centering and cropping operation to standardize all images to a uniform 200×950 pixel dimensional image. (b) Diagram illustrates how zebrafish in each clutch is re-labeled before cLDM training, in order to prevent label leakage during downstream classification. There are two example clutches in this illustration, named by the date of collection: 2022.11.11 and 2022.11.08. Clutch 2022.11.11 profiles individuals with the arhgap11A and control genotypes, and clutch 2022.11.08 profiles individuals with the fam72B and control genotypes. After relabeling, the arhgap11A genotype individuals are given the label arhgap11_1, which is unique and not shared with arhgap11A genotype individuals from other clutches. For the controls, each clutch’s controls are divided into two groups: one with a common label across all clutches (“Controls”), and clutch-specific controls (“hoControls_1”, “hoControls_2”).

After relabeling all training images, we run cLDM analysis on the entire image set. After training, we identify the clutch with the largest set of controls, and define that clutch as the reference clutch. We then pass all training images through the trained cLDM, but during reconstruction we specify the reference clutch during the reverse denoising process, so that all images are reconstructed in the style of the reference clutch (**Figure 1c**). The adjusted images are then used for unsupervised clustering analysis and downstream classification, as discussed below.

### cLDM mitigates clutch effects in zebrafish imaging

To evaluate the ability of cLDM to remove the clutch and age effect from the zebrafish images, we trained cLDM on all 3,146 zebrafish images, using the labeling scheme discussed in the previous section. To be clear, in the cLDM input, only the common control individuals have shared genotype labels across clutches, and every combination of genotype, clutch and age group are given their own unique label. As a result, cLDM explicitly treats individual zebrafish of the same genotype (but from different clutches or ages) as different classes; thus, any similarity that arises between zebrafish from the same genotype but different clutches after cLDM processing will be unsupervised. For downstream analysis, after training, we reconstructed every training image using a selected reference clutch during reconstruction such that all training images were stylized in a single reference clutch, and used the image embeddings (without reconstruction by the VAE decoder) for analysis.

As comparison, we applied the Safe Self-Refinement for Transformer-based (SSRT) Domain Adaptation method to the same dataset. SSRT is a transformer-based domain adaptation method that employs a safe self-refinement strategy to iteratively update the model and reduce errors during domain adaptation^14^. The SSRT method requires samples in both a source and target domain, which we constructed as follows. For each mutant genotype, we selected one clutch and placed it in the source domain, while all other clutches containing the same genotype label were assigned to the target domain. For controls, the common controls were all placed in the source domain, while all held-out controls were placed in the target domain (**Figure 3a**). To make the results comparable to those of cLDM, we applied SSRT to the VAE-embedded zebrafish images that cLDM were trained on as well.

**Figure 3:**
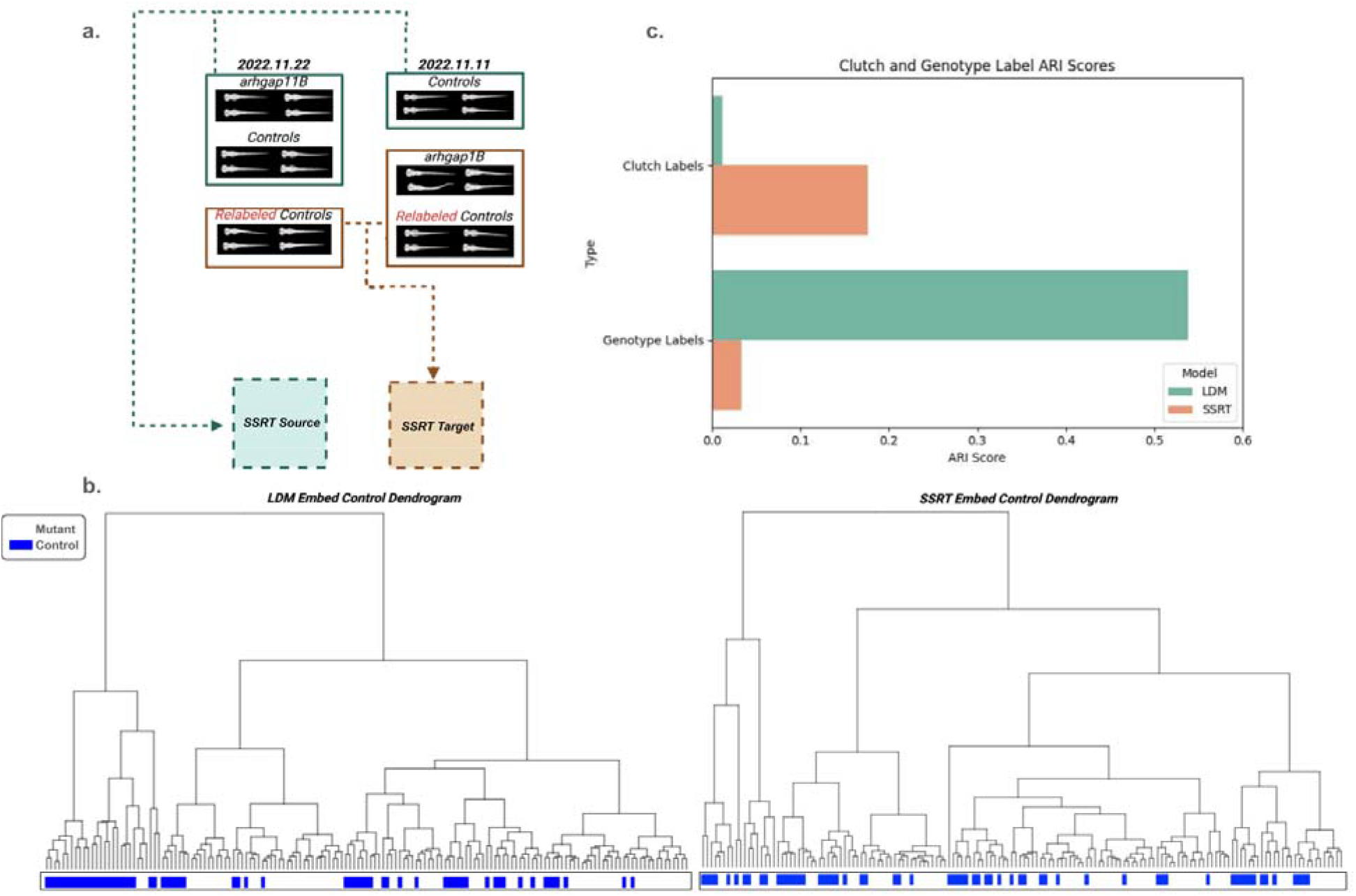
Benchmarking the clutch mitigation performance of cLDM and domain adaptation methods on the zebrafish data (embedding space). **(a)** Illustration of how data is split into the source and target domain for training for SSRT. (b) Dendrograms illustrating the clustering of zebrafish images in embedding space after processing by cLDM (left) or SSRT (right). Blue indicates held-out control label groups, white indicates held-out mutant label groups. **(c)** Quantitative performance of cLDM and SSRT on clutch mitigation. Performance is evaluated using two ground truth labels: clutch labels and genotype labels. Lower clutch ARI indicates better performance (indicating zebrafish are not clustering by clutch). Higher genotype ARI indicates better performance (indicating zebrafish with the same genotype but profiled on different clutches are still clustering based on image embedding).

After training cLDM and SSRT on all zebrafish images, we first clustered the reconstructed image embeddings using hierarchical clustering separately for each method using Fréchet Inception Distance (FID)^6^ as a distance metric (**Fig. 3b**). It is clear that the held-out control genotype groups cluster more strongly from cLDM processing compared to SSRT. This behavior is unexpected because during cLDM processing, each of the held-out genotype groups was given a unique genotype label, so the fact they cluster more strongly after cLDM processing suggests that cLDM is more strongly able to remove the clutch effect compared to SSRT. **Figure 3c** quantifies the degree to which cLDM and SSRT are able to remove the clutch effect. More specifically, for each of the held-out genotypes, we have its ground truth label with respect to which clutch it belongs to, as well as whether the genotype refers to a mutant or control genotype. Effective clutch removal should yield lower consistency, as measured by the adjusted rand index (ARI), between a method and the ground truth clutch labels, because when clutch effect is removed, then genotypes should not group by clutch. Similarly, better clutch removal should yield higher consistency between the method and the ground truth mutant labels (indicating the held-out control groups cluster). We see cLDM outperforms SSRT on both the clutch and mutant label groupings, achieving a 93.6% reduction in ARI for clutch grouping (0.01 compared to 0.17), and 98.19% increase in ARI for mutant grouping (0.03 compared to 0.53) (**Fig. 3c**).

### cLDM clutch effect mitigation also evident in image space

In the benchmarking we performed above, clutch effect removal comparison was performed in image embedding space to ensure the results were not driven by noise introduced by the VAE decoder during image reconstruction. However, the final goal of our analysis is to identify genotype-specific morphological effects, which require analysis at the image level, not the embeddings. We therefore aimed to check cLDM-reconstructed images to ensure they are also consistent with mitigated clutch effects.

We took the image embeddings after cLDM processing from the previous section, and reconstructed them into image space using the trained VAE decoder. We then compared cLDM-reconstructed images of different genotypes and clutches by performing hierarchical clustering and computing FID distances, and comparing the results to clustering based on the original images (**Fig. 4**). Looking at the original images, the zebrafish overall cluster by age (3dpf vs 5dpf), but have less evident structure beyond that (**Fig. 4a**). Notably, with respect to the original images, the control images do not group together, consistent with the presence of strong clutch effect. In contrast, cLDM-processed images principally cluster by controls as one smaller cluster, and mutants as another cluster (**Fig. 4b**). Our results suggest that cLDM correction also was effective at mitigating clutch effects in the image space.

**Figure 4:**
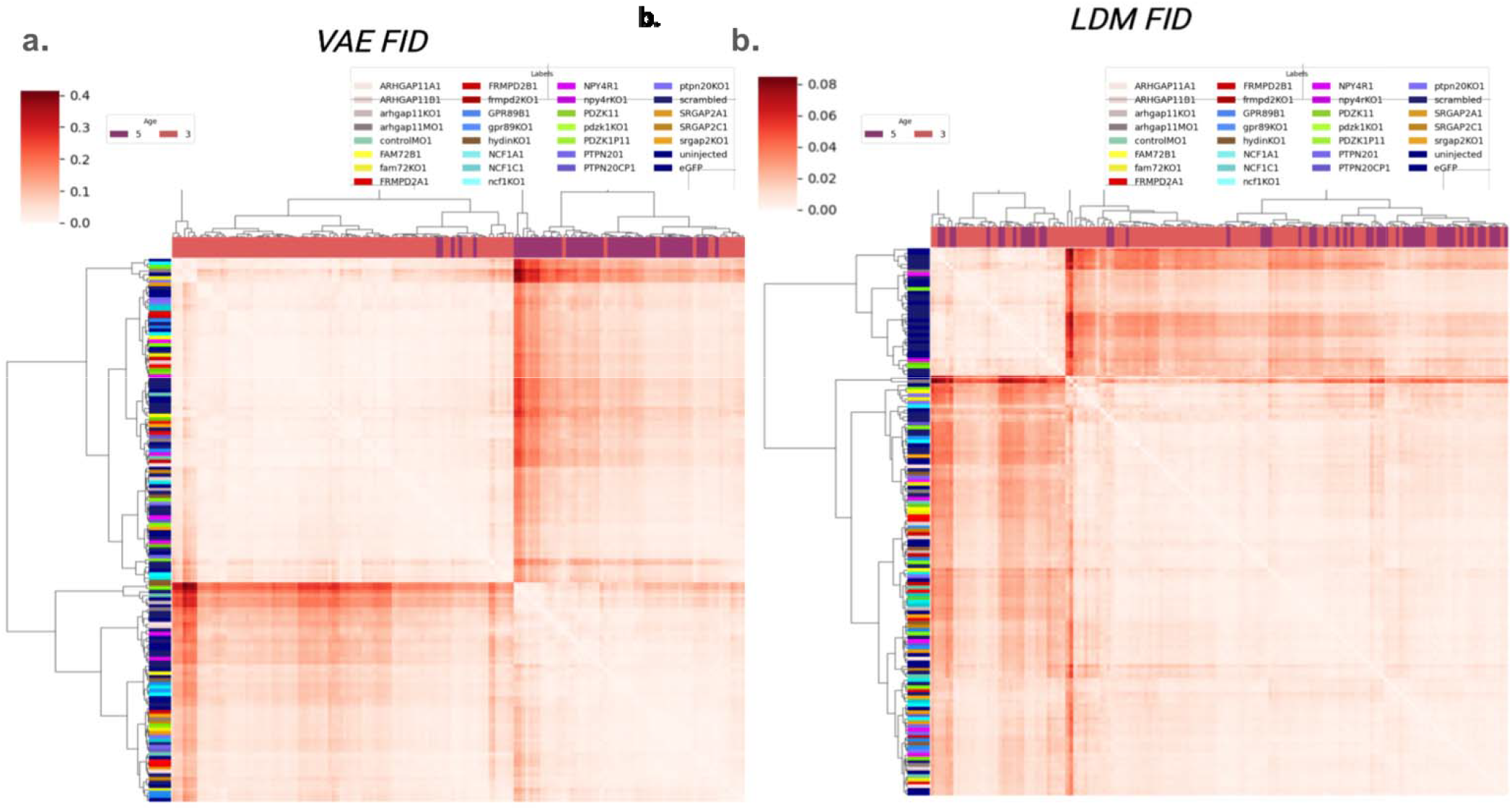
Clustering of cLDM-adjusted zebrafish images compared to VAE only-adjusted images. **(a)** To determine the extent to which zebrafish with similar genotype but profiled from different clutches cluster, we passed the original zebrafish images through the encoder and decoder of the VAE (without the cLDM), and clustered. Each row and column correspond to a unique combination of clutch and mutant genotype label. Columns are labeled by the age of the zebrafish. Rows are labeled according to genotype. Heatmap indicates FID between pairs of zebrafish groups. **(b)** Same as (a), but FID calculations were made based on the original zebrafish images that were passed through the VAE encoder, cLDM forward and reverse process (conditioned on a single reference clutch), and VAE decoder.

### Mutant genotypes form two distinct morphologies

Having confirmed the clutch effect is mitigated in the adjusted zebrafish images, we next explored how the adjusted images might reveal morphological similarities across the genotypes. A longstanding goal of molecular biology is to identify genotype-phenotype relationships, that is, to understand how genetic perturbations yield phenotypic (or in this case, morphological) variation. When two similar genetic perturbations yield similar morphological changes, it suggests those two perturbations might be acting on a similar molecular pathway, yielding functional insight into the genome. Our goal then was to determine whether different mutants (profiled across clutches) exhibited similar impacts on morphology, which would suggest shared molecular pathways. To this end, we designed a series of binary classification experiments in which for every pair of genotypes, we fine-tuned an AlexNet classifier^15^ to distinguish the two genotypes, and recorded how often zebrafish of one genotype was either correctly classified, or mis-classified as the other genotype. We interpret poor classification performance as morphological similarity between two genotypes: if two genotypes yield similar morphologies, it will be difficult to distinguish their images.

To train our binary classifiers, we implemented a two-stage training procedure due to the small size of the zebrafish dataset. We first subsetted the original dataset into 10 family datasets, where all genotypes corresponding to a single gene family was combined with all controls to form a single family dataset (see Supplementary Tables for assignment of genotypes to family). For each family dataset, we fine-tuned an AlexNet model first on each family dataset, then further fine-tuned the family-trained dataset on each genotype’s data specifically to train the binary classifier. We chose AlexNet due to its smaller architecture and ease of training, which were important since we trained 450 models. To measure classification performance, we computed Area Under the Receiver Operating Characteristic curve (AUROC) scores between all pairs of mutants.

**Figure 5** illustrates all binary classification performance estimates for all pairs of genotypes, using either the original images (**Fig. 5a,b**), or the cLDM-adjusted images (**Fig. 5c,d**). When trained using the original images, there is no clear relationship between genotypes; it is only mildly apparent that the controls are systematically somewhat different from the other genotypes (average AUROC=0.64). In contrast, when trained on the adjusted images from the cLDM, the a number of key trends become apparent (**Fig. 5c,d**). First, the non-control genotypes broadly form two groups of mutants, with the GPR89 family (grp89KO, GPR89B), FRMPD family (FRMPD2A, FRMPD2B, frmpd2KO), FAM72 family (FAM72B, fam72KO), NCF family (ncf1KO, NCF1A, NCF1C) and SRGAP2 family (SRGAP2A, SRGAP2C, srgap2KO) forming one group, while the NPY4R family (npy4rKO, NPY4R), PTPN20 family (PTPN20, PTPN20CP, ptpn20KO), PDZK1 (pdzk1KO, PDZK1P1, PDZK1), and ARHGAP11 family (arhgap11MO, arhgap11KO, ARHGAP11A, ARHGAP11B) forming another group, suggesting broad similarities in the morphological effects within each group. Furthermore, the controls are more distinct from the mutants in the cLDM-adjusted images (AUROC = 0.96) compared to the original images (AUROC= 0.64), suggesting better detection of genotype effects.

**Figure 5:**
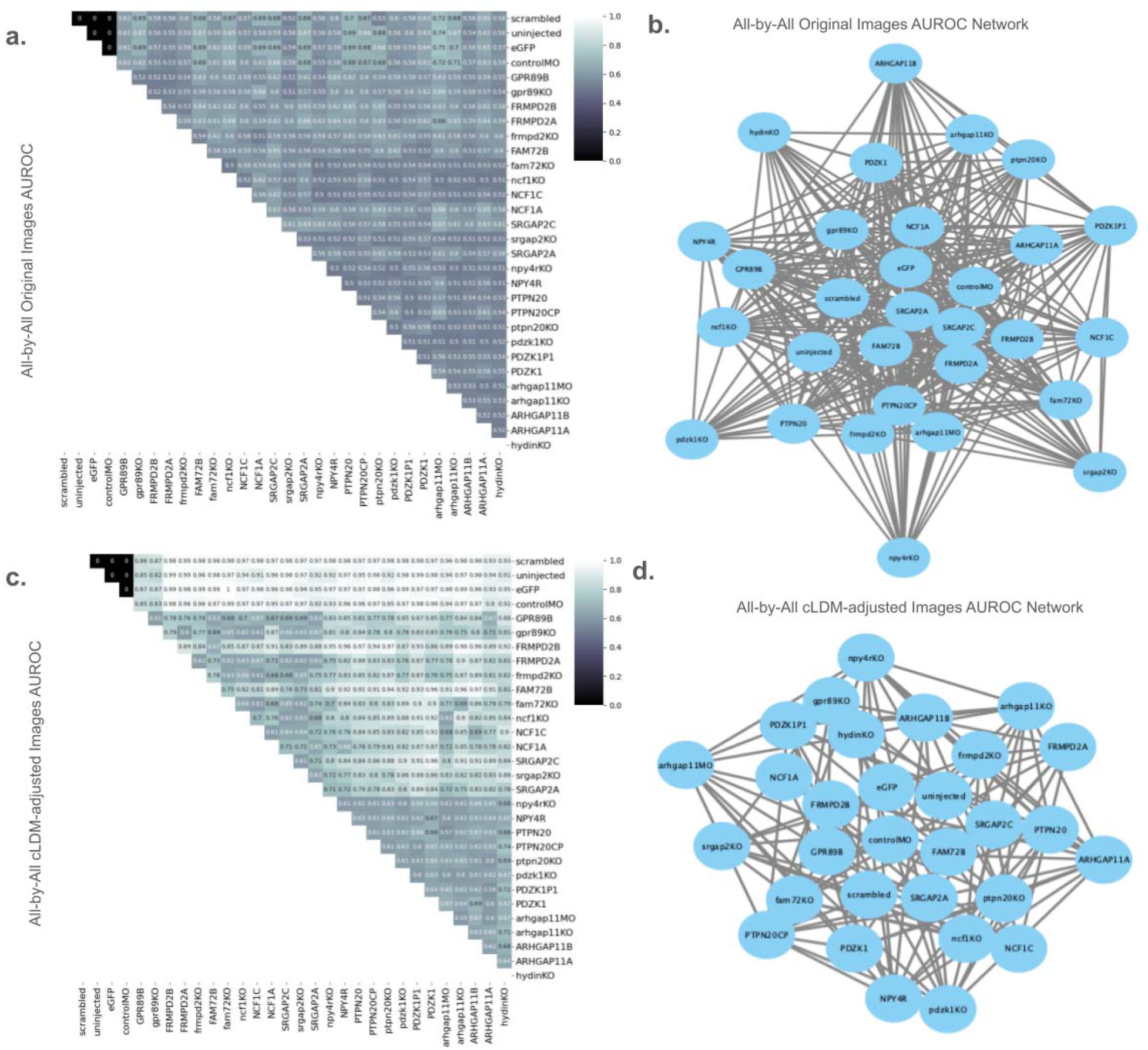
Genotypes form two morphologically distinct clusters, separated from controls. **(a)** Performance of binary classifiers trained to distinguish pairs of genotypes using original zebrafish images. Performance is measured as AUROC. (**b**) Network representation of (a), where nodes represent genotypes, and edges represent high misclassification rate between the two genotypes, suggesting that they have similar morphology (AUROC < 0.6). (**c**) Same as (a), but training images are the cLDM-adjusted images. (**d**) Same as (b), but with respect to the classifiers trained in (c).

Our results closely align with previously validated experimental findings. For instance, we observed similar patterns for two mutant models of *GPR89*, which achieved the highest F1 scores in our classification task. Feature attribution methods further highlighted this gene, showing the highest gradient values concentrated in the head region. These findings align with prior results that identified *GPR89* as having one of the strongest morphological signals among all of the mutant models, particularly with a significant effects on the zebrafish larval head and brain size^4^. Conversely, the *PDZK1* mutant models demonstrated the lowest scores in our classification task and, consistent with experimental findings, exhibited the most morphological similarity to their associated controls compared to other mutant families. Taken together, these results suggest that the cLDM model not only mitigates clutch effects but also enhances the model’s ability to uncover biologically meaningful morphological similarities among mutants.

## DISCUSSION

Studies of living organisms often involve measuring how data modalities such as imaging vary as a function of multiple complex factors related to biology (genetics, the environment, age, conditions) and technical confounders (such as batch). When the number of individuals per batch is small relative to the dimensionality of the biological factors, it can be challenging to remove batch effects from data modalities such as imaging. While specialized domain adaptation techniques such as multi-source domain adaptation and universal domain adaptation are designed to handle classification problems where not all classes are present in every batch, they cannot handle cases where there are more than one label sets (e.g. beyond class labels, there are other biological factors that must be accommodated across batches, such as age and genotype in this study). Here we show that a conditional latent diffusion model is capable of addressing batch correction when each individual data point has multiple label sets that must be addressed. Our training approach provides a favorable framework for these types of problems and represents a novel application for addressing such imaging scenarios.

Imaging problems such as that addressed in this paper arise frequently in medicine (from multimodal data analysis problems involving human medical records and medical diagnostic imaging) and biology (where model organisms are grown in clutches/litters and specific biological variables manipulated) where rich metadata on individuals is available. This is unlike in many problems in computer vision, where it can be possible to obtain large datasets of unlabeled data, and where the traditional assumptions of domain adaptation methods may be more suited. Thus, we expect our approach to domain adaptation to be most suited in the life sciences and medicine.

Through our analysis, we demonstrate that successfully addressing the clutch effect can have striking impact on the conclusions drawn from the data at hand. More specifically, without addressing the clutch effect, little to no structure is evident from the genotype classification experiment, other than differences between each mutant genotype versus control. However, after clutch mitigation, it becomes clear that the mutant genotypes form two groups of distinct morphologies that are also distinct from controls, suggesting that the mutant genotypes in total target two phenotypically-relevant pathways. Compared to a non-neural network-based analysis of the same data that focused on comparing hand-engineered imaging features, we also detect significantly more morphological differences and similarities between genotypes, suggesting our model can enrich data analysis of imaging data with complex metadata. ^4^

## ACKNOWLEDGEMENTS

This work was supported, in part, by U.S. National Institutes of Health (NIH) grants from the Office of the Director and National Institute of Mental Health (R01MH132818 to M.Y.D., DP2MH129987 to G.Q), National Institute of Child Health and Human Development (P50HD103526 to G.Q). This work was supported, in part, by NSF CAREER award (1846559 to G.Q.).

## METHODS

### cLDM overall architecture

We adopted the framework of the Latent Diffusion Model (LDM) from Rombach et al^16^., in which image embeddings are passed through a forward diffusion process and reverse denoising process (**Fig. 1**). The forward process incrementally adds Gaussian noise to the embeddings over a fixed number of time steps, while the reverse process, modeled by a U-Net, iteratively denoises embeddings such that after the final denoising step, image embeddings approximate the original input embedding. Each step of the reverse denoising process is conditioned on biological and technical factors of interest (genotype, clutch, age) to encourage cLDM to learn clutch, age and genotype-specific effects on imaging.

### Variational Autoencoder

To generate compact latent representations for the cLDM, we trained a convolutional Variational Autoencoder (VAE). The encoder compresses each image *x*_*i*_ into a latent representation *z*_*i*_. It consists of convolutional layers that reduce spatial resolution while increasing feature depth, outputting the mean *µ* (*x*_*i*_) and log-variance *log log σ*^2^ (*x*_*i*_) of a diagonal Gaussian posterior, of dimension 2304 (48 × 48). A sample is drawn from the Gaussian posteriorThis approximates to a 48×48 latent grid which is used as the input of the LDM. The latent vectors are sampled using the reparameterization trick:

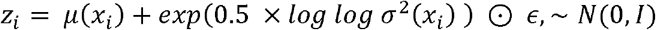

The sample is reshaped into a 48×48 latent grid which is used as the input to the cLDM.The decoder mirrors the encoder’s structure using transposed convolutions to reconstruct the embeddings back into the original image space. A final layer with tanh activation produces the output image 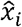 within the original intensity range. The model is optimized using a loss that combines pixelwise mean squared error (MSE) with the Kullback–Leibler divergence (KL) between the approximate posterior and the standard Gaussian prior:

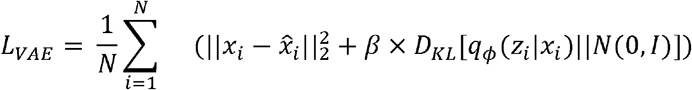

In our implementation the VAE’s role is that of a perceptual compression module, used to obtain informative latent representations of the input images. The encoder consists of four convolutional layers: two initial resizing layers with strides (1,2), followed by two stride convolutions that progressively down sample the input to a final shape of (48 × 48). Batch normalization and LeakyReLU are applied throughout. The decoder mirrors this structure using five transposed convolutional layers to up sample the latent representation back to the original image resolution. In total, the encoder and decoder process approximately 3.1 million and 1.8 million spatial nodes, respectively. We further assign a small weight value to the KL divergence weight *β* prioritizing accurate reconstruction rather than enforcing strict regularization of the latent space. All weights in linear layers are initialized using Xavier initialization^17^, and batch normalization^18^ is used throughout for stable training.

### Conditional Diffusion Model

The diffusion model (DM) relies on the latent embeddings *z*_*i*_ which are obtained from the encoder for each image *x*_*i*_. Here we refer to *z*_*i*,0_ as the latent representation of the original image *x*_*i*_ image at timestep 0 and use *z*_*i,t*_ to denote the embedding at diffusion timestep *t*, where *t* ∈ {1,…, *T*} in which *T* = 350 by default in this study. The forward process of the DM adds Gaussian noise to *z*_*i*,0_ through a predefined linear noise scheduler, controlling the rate at which noise is injected at each time step over *T* until reaching *z*_*i,t*_ this is calculated as:

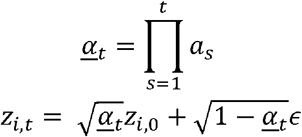

Where *<u>α</u>*_*t*_ is the cumulative product of the noise scheduler, and *ϵ ∼ N* (0,1), is standard Gaussian noise. This ultimately transforms the initial *z*_*i*,0_ into a noised version *z*_*i,T*_.

In the reverse process, we reconstruct the *z*_*i*,0_ by iteratively denoising the *z*_*i,T*_. It’s during this step in which we can introduce a condition variable *c* to guide the output to a desired batch. In our implementation, the condition *c* corresponds to the batch variable *b*, allowing the model to generate outputs consistent with the desired experimental batch. This conditional reverse process can be expressed as:

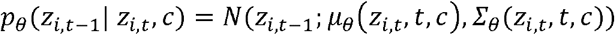

Where *µ*_*θ*_ and *Σ*_*θ*_ represent the predicted mean and variance of the denoised distribution. The *ϵ*_*θ*_(*z*_*i,t*_,*t,c*) denoising model follows a U-Net-like architecture adapted for latent inputs, consisting of down sampling and up sampling blocks with residual connections, group normalization, timestep embeddings. Ultimately the reconstructed embedding *z*_*i*,0_ is decoded through the VAE decoder to produce the output image *x*’, which represents the original image *x*_*i*_ in the style of our desired batch.

Our LDM is trained by minimizing the mean squared error between the true *ε* and predict noise *ε*_*θ*_(*z*_*i,t*_,*t*) expressed as:

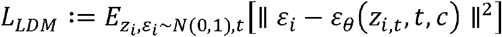

This ensures that the model is accurately predicting the noise applied during the forward process, and by conditioning on the batch, we ensure that the model can reconstruct the morphological images in the style of desired batch with respect to the original mutant.

We trained the model using a linear noise scheduler for 2000 epochs, early stopping based on the best validation loss (patience = 10) and a learning rate of 1×10−4. The loss was computed as the mean squared error (MSE) between the ground truth noise and the predicted noise. Based on preliminary experiments, we determined that using 350 noise steps in the forward diffusion process provided the optimal balance between reconstruction fidelity and sampling efficiency.

### Construction of zebrafish conditioning variables

Each sample dataset is associated with an age, a batch identifier, and genotype label which is either a control or mutant. To encourage the model to learn the batch effect, rather than features associated with age or genotype, we isolated the batch variable as the only consistent variable. This was achieved by creating a unique encoding for each combination of mutant/control status and batch identity to remove any shared mutant or control labels across batches, thus preventing the model from inferring identity through genotype. We then subset each control batch into two groups, applying a unique encoding to one group while retaining the original encoding for the other half; this enables the model to learn the relationship between the control batches. After generating the new encodings, we applied an 80/20 split to create the training and validation sets, which were used for early stopping during model training.

### cLDM inference process

After training cLDM on the entire zebrafish imaging data, we perform inference to reconstruct new zebrafish images, adjusted for clutch effect. For a novel input image *x*_*input*_, we first apply the frozen VAE encoder to obtain its latent representation *z*_*rep*_. To guide the reconstruction toward a specific batch, we introduce a conditioning label c, which encodes the target batch identity. The latent representation *z*_*rep*_ is then passed through the forward diffusion process, where Gaussian noise is progressively added over 350 timesteps to produce a noisy embedding *z*_*noisy*_.

In the reverse process, the trained U-Net denoises *z*_*noisy*_ back to a new latent *z*_*rep*_ conditioned on *c*. This conditional denoising allows the model to reconstruct an image as though it were drawn from the distribution of the desired batch or consistent with the target batch. Finally, the denoised representation is passed through the frozen VAE decoder to yield the final output image *x*’, representing the input sample rendered in the style of the target batch.

### Assessment of Batch Normalization and Mutant Separability

To evaluate the model’s ability to correct for batch effects, we designed a series of experiments involving the computation of FID and AUROC scores. First, all samples were passed through the LDMZ while conditioning on the largest control plate, thereby perturbing each sample into a common batch. The resulting images formed the perturbed dataset, which we used for downstream analysis. The original, unaltered images were retained as the unperturbed dataset for baseline comparison.

### Clustering of zebrafish via FID Clustering

To assess group-level similarity, we computed pairwise Fréchet Inception Distance (FID) scores between all groups in the perturbed dataset, defined as:

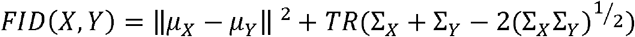

Here *µ*_*X*_ and *µ*_*Y*_ are the mean feature vectors while *Σ*_*X*_ and *Σ*_*Y*_ are the covariance matrices of the two distributions being compared. The first term measures the squared difference between the means, while the trace term quantifies differences in the covariance structure. These scores were visualized using Seaborn’s clustermap function to reveal the structure of inter-group similarity. The same procedure was applied to the unperturbed dataset to enable comparison.

### Binary classification experiments

To assess the discriminability between all individual samples, we conducted an all-by-all classification experiment meaning we systematically compared every pairwise combination of classes (e.g., mutants and controls) using binary classification. For each pair of classes (e.g., mutant A vs. mutant B, mutant A vs. control, mutant B vs. control, etc.), we fine-tuned a pre-trained AlexNet model^15^. Specifically, we replaced the final fully connected classification layer to perform binary classification and used a relatively low learning rate. Our goal was to distinguish between just those two classes, and computed AUROC scores. This process was repeated across all unique pairs across both the unperturbed and perturbed datasets.

### Data Preprocessing

#### Image acquisition and raw quality control

Details of experiments related to generation of mutant and control larvae and imaging can be found in Soto et al^4^. Briefly, a total of 3,146 zebrafish larvae were imaged at 3 or 5 days post fertilization using a vertebrate automated screening technology (VAST)^19^ at a resolution of 200×1200 pixels with 3-bit depth. The LP Sampler anaesthetizes individual zebrafish larva positioned in a well of a 96-well plate and through a capillary automatically images each larvae from four perspectives (anterior lateral, posterior lateral, ventral, and dorsal).^20^ Any damaged or dead fish or truncated fish that were not fully visible in the well were flagged as issues in the original collection and were removed in the initial stages of preprocessing.

#### Segmentation and background removal

After initial quality control,we generated segmentation masks for each sample to accurately remove the background for all the images. This was achieved through the use of the Segment Anything Model (SAM) version 1 and the and the FishInspector imaging software v1.03 (Figure 4a).^21^ FishInspector generated JSON files with centroid pixel coordinates for different landmark features; we extracted the coordinates for three features (yolk, contour, tail) and passed those to the SAM model (Figure 5a). The model generated segmentation masks for each image based on the three landmark features. We set all of the pixels outside of the generated mask to black. To address some images which cut off the zebrafish, we then removed any samples whose segmentation mask overlapped with the right-most or left-most edges of the image within a 5-pixel wide boundary (**Figure 5a**).

#### Spatial normalization

All retained images were subsequently processed further by centering, cropping and padding to a shape of 200×794 pixels. Images were centered in the 200×1024 pixel space and cropped down to their smallest possible bounding box. Padding was applied by adding black pixels to resize the images to a uniform shape of 200×794 pixels. To further differentiate the zebrafish from the background, we inverted the pixel values for each sample, retaining the fully black background. The resulting curated dataset comprised 3450 centered images.

## Code availability

The implementation of cLDM can be found at https://github.com/quon-titative-biology/cLDM.

## Notes

https://github.com/quon-titative-biology/cLDM

### Competing Interest Statement

The authors have declared no competing interest.

